# Mechanistic insights into ventricular arrhythmogenesis of hydroxychloroquine and azithromycin for the treatment of COVID-19

**DOI:** 10.1101/2020.05.21.108605

**Authors:** Gongxin Wang, Chieh-Ju Lu, Andrew W. Trafford, Xiaohui Tian, Hannali M Flores, Piotr Maj, Kevin Zhang, Yanhong Niu, Luxi Wang, Yimei Du, Xinying Ji, Yanfang Xu, Lin Wu, Dan Li, Neil Herring, David Paterson, Christopher L.-H. Huang, Henggui Zhang, Ming Lei, Guoliang Hao

## Abstract

**Aims:** We investigate mechanisms for potential pro-arrhythmic effects of hydroxychloroquine (HCQ) alone, or combined with azithromycin (AZM), in Covid-19 management supplementing the limited available experimental cardiac safety data.

**Methods:** We integrated patch-clamp studies utilizing In Vitro ProArrhythmia Assay (CiPA) Schema IC_50_ paradigms, molecular modelling, cardiac multi-electrode array and voltage (RH237) mapping, ECG studies, and Ca^2+^ (Rhod-2 AM) mapping in isolated Langendorff-perfused guinea-pig hearts with human in-silico ion current modelling.

**Results:** HCQ blocked I_Kr_ and I_K1_ with IC_50_s (10±0.6 and 34±5.0 μM) within clinical therapeutic ranges, I_Na_ and I_CaL_ at higher IC_50_s, leaving I_to_ and I_Ks_ unaffected. AZM produced minor inhibition of I_Na_, I_CaL_, I_Ks_, and I_Kr_,, sparing I_K1_ and I_to_. HCQ+AZM combined inhibited I_Kr_ and I_K1_ with IC_50_s of 7.7±0.8 μM and 30.4±3.0 μM, sparing I_Na_, I_CaL_ and I_to_. Molecular modelling confirmed potential HCQ binding to hERG. HCQ slowed heart rate and ventricular conduction. It prolonged PR, QRS and QT intervals, and caused prolonged, more heterogeneous, action potential durations and intracellular Ca^2+^ transients. These effects were accentuated with combined HCQ+AZM treatment, which then elicited electrical alternans, re-entrant circuits and wave break. Modelling studies attributed these to integrated HCQ and AZM actions reducing I_Kr_ and I_K1_, thence altering cell Ca^2+^ homeostasis.

**Conclusions:** Combined HCQ+AZM treatment exerts pro-arrhythmic ventricular events by synergetically inhibiting I_Kr_, I_Ks_ with resulting effects on cellular Ca^2+^ signalling, and action potential propagation and duration. These findings provide an electrophysiological basis for recent FDA cardiac safety guidelines cautioning against combining HCQ/AZM when treating Covid-19.

## INTRODUCTION

The severe acute respiratory syndrome coronavirus 2 (SARS-CoV-2) pandemic has now infected >5.18 million individuals causing 338039 deaths as of 23-May-2020 (www.ecdc.eurpoe.eu). Although there are no approved drugs to prevent or treat SARS-CoV-2 infection, recent use of the anti-malarial 4-aminoquinolines chloroquine (CQ) or hydroxychloroquine (HCQ) alone, or combined with the antibiotic azithromycin (AZM) in Covid-19 therapy has attracted global attention and extensive early clinical use and trials(1, 2). An initial nonrandomized open label clinical trial of 20 cases, had associated HCQ treatment with viral load reduction or disappearance in COVID-19 patients, effects reinforced by AZM(3). This subsequently prompted the US Food and Drugs Administration (FDA) authorization of emergency use of CQ or HCQ, alone or in combination with AZM in Covid-19 therapy(4, 5). Currently, large number of trials of these combinations, with or without the addition of further drugs, are registered worldwide (1, 2).

However, several recent studies have flagged cardiac safety concerns for the CQ/HCQ and AZM regime in Covid-19 therapy. Particular concern has arisen regarding potential synergistic effects of HCQ and AZM on QT duration and cardiovascular mortality (6–10). Higher mortalities and incidences of prolonged corrected QT (QTc) intervals were reported in patients at a Brazilian hospital receiving the high dose (12 g total dose over 10 days, ~1.2 g/day, giving plasma levels of ~10 μM) than the lower dose of CQ, typified by its use in other conditions such as rheumatoid arthritis and systemic lupus erythematosus (SLE). This phase 2b randomized Covid-19 trial was terminated after enrolment of only 81 out of the target of 440 patients owing to occurrences of adverse events (6). A retrospective data analysis from United States Veterans Health Administration medical centers suggested increased overall mortality even in patients treated with HCQ alone (7). The largest available analysis from an international consortium on HCQ safety covering over 950,000 patients from six countries declared HCQ safe for short-term use at doses currently used for conditions such as rheumatoid arthritis. However, it urged caution in its use when combined with AZM in view of potential risks of cardiovascular mortality related to QTc prolongation(8). Two recent reports of 90 and 201 patients hospitalized with COVID-19 treated with HCQ, with or without AZM, associated these drugs with QTc prolongation, especially when used in combination(9, 11). Although these studies are small, only one case of torsades de pointes was documented and the therapy was seldom discontinued. They suggested that more QT interval monitoring studies should be undertaken before final recommendations can be made(11). Based on these findings, the FDA issued a cautions warning against use of HCQ or CQ for COVID-19 outside of the hospital setting or a clinical trial, due to risks of heart rhythm problems(4). This does not affect FDA-approved uses in malaria, systemic lupus erythematosus (SLE), and rheumatoid arthritis where they have been used many years. Both HCQ and AZM have been in clinical use alone or in combination in other disease conditions for many years with cardiac safety then not considered a major concern.

Currently, there is very limited data evaluating cardiac safety particularly concerning use of HCQ in combination with AZM, although multiple randomized trials are currently in progress evaluating the efficacy of this combination for Covid-19. We therefore explored the cardiac safety of HCQ alone or in combination with AZM using the FDA recommended Comprehensive in Vitro Proarrhythmia Assay (CiPA) protocols to understand arrhythmic mechanisms (12). Furthermore, we expanded the CiPA protocol by introducing assessments of effects on membrane and Ca^2+^ clocks, their interaction and cardiac conduction recently suggested as crucial indicators for ventricular arrhythmogenesis (13, 14).

## RESULTS

The four groups of experiments conducted assessed the mechanistic basis for the clinical reports bearing upon cardiac safety of HCQ, whether applied alone or in combination with AZM.

### Patch-clamp studies of single channel characteristics

The first series of experiments assessed the effects of HCQ (Fig. 1) and AZM alone (Fig. 2) and in combination (Fig. 3) on a range of human cardiac ion channels. The experiments on HCQ and AZM themselves adopted concentrations fully encompassing plasma levels associated with their clinical use. The same HCQ concentration range was then combined with 10 μM AZM. These examined inward, cardiac channel currents from hNav1.5 (I_Na_, *A*), Cav1.2 (I_Ca,L_, *B*), mediating or maintaining membrane action potential depolarization. They also studied outward Kv4.3 (I_to_, *C*), hERG (I_Kr_, *C*), KCNQ1/E1 (I_Ks_, *E*) and Kir2.1 (I_K1_, *F*) normally mediating action potential recovery. This encompassed the range of ion channels recommended by the CiPA protocol.

**Figure 1.**
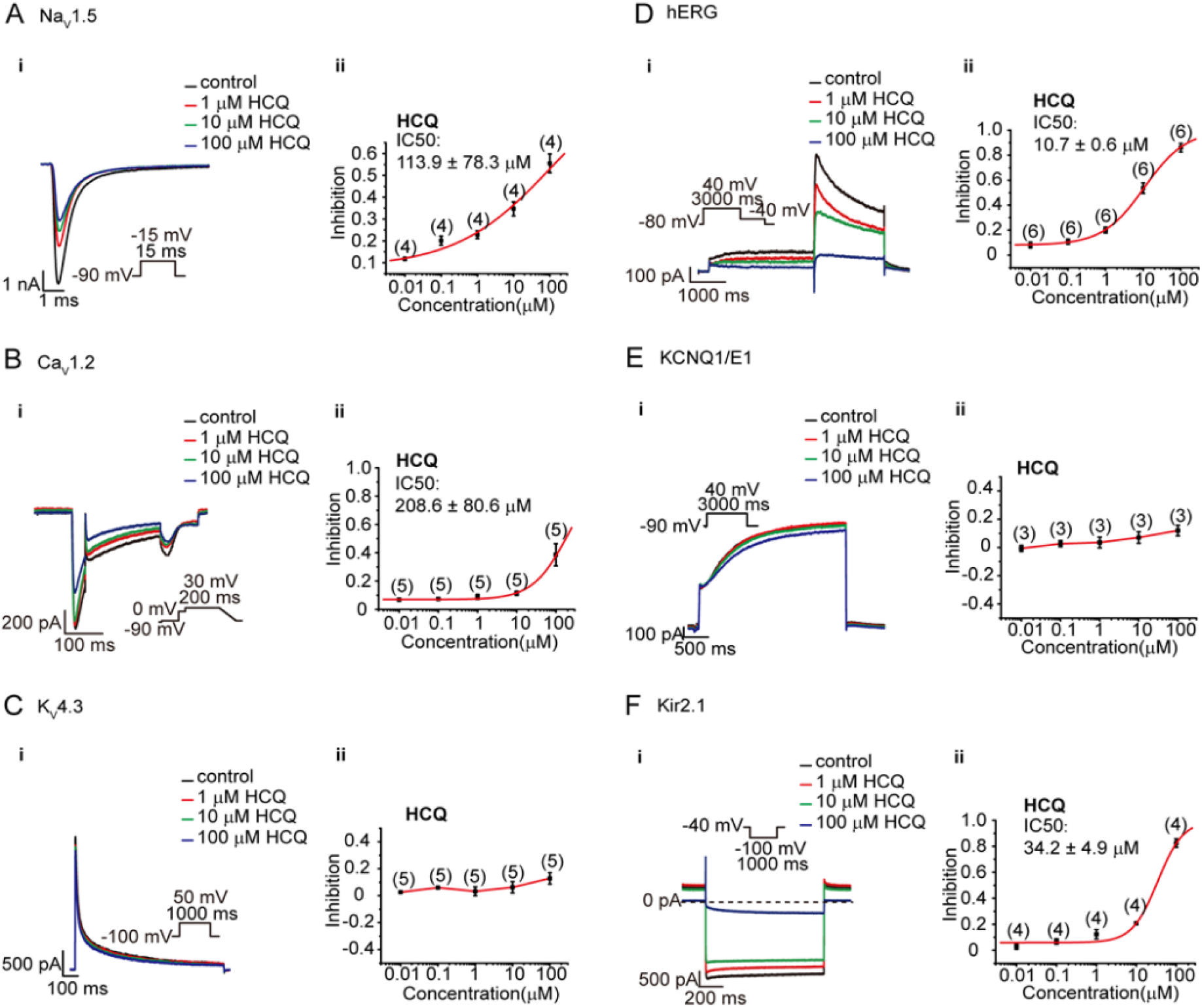
Effects of HCQ on ion currents attributable to hNav1.5 (I_Na_, A), Cav1.2 (I_Ca,L_, B), Kv4.3 (I_to_, C), hERG (I_Kr_, D), KCNQ1/E1 (I_Ks_, E) and Kir2.1 (I_K1_, F). (A-F) Left panels (i): Typical currents in the presence of 0, 1, 10 and 100 μM HCQ. Insets summarize the voltage clamping protocols. Right panels (ii): Mean concentration-response plots constructed for the maximum magnitude of each current (n cells).

**Figure 2.**
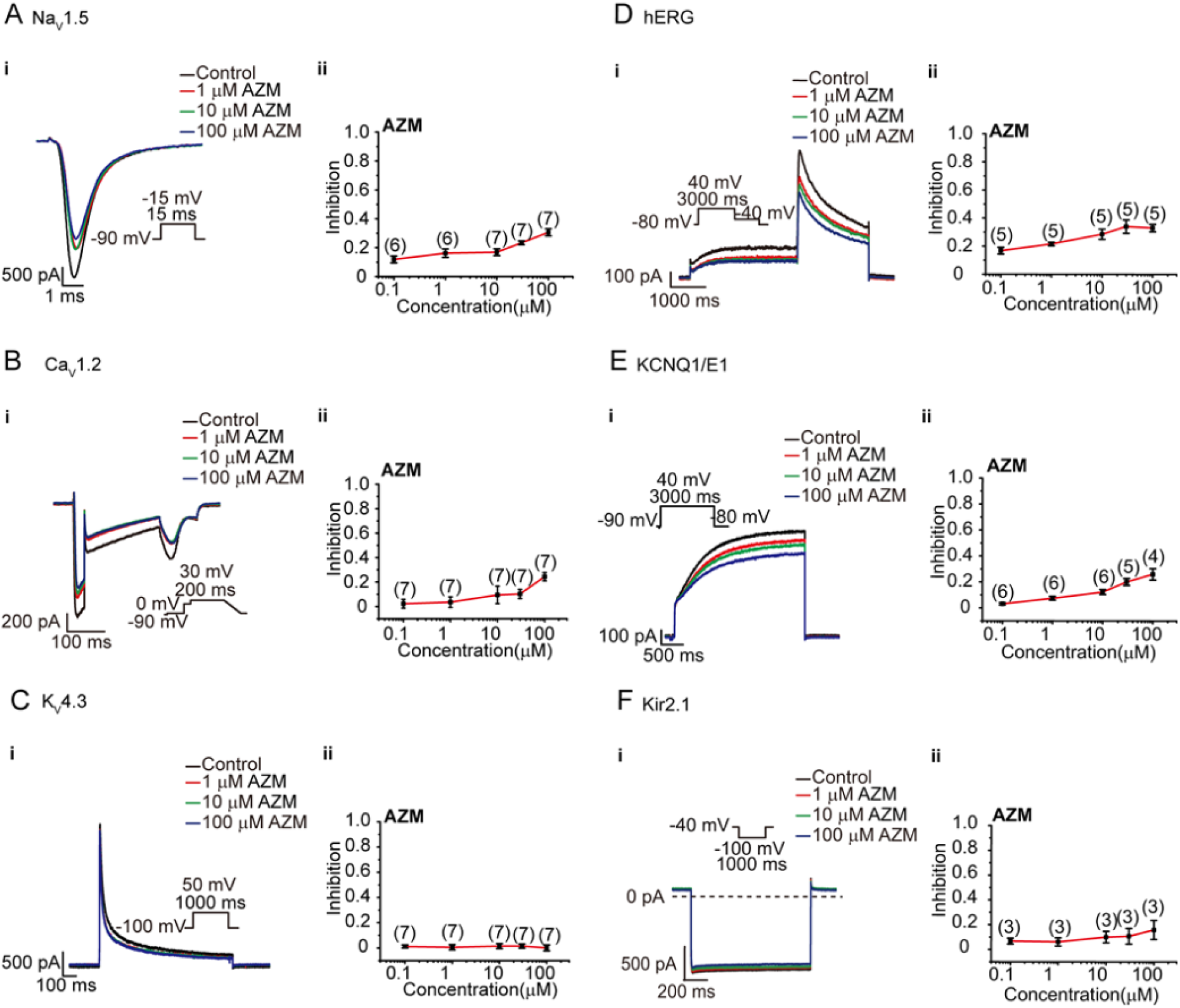
Effects of AZM on ion currents attributable to hNav1.5 (I_Na_, A), Cav1.2 (I_Ca,L_, B), Kv4.3 (I_to_, C), hERG (I_Kr_, *D*), KCNQ1/E1 (I_Ks_, E) and Kir2.1 (I_K1_, F). (A-F) Left panels (i): Typical currents in the presence of 0, 1, 10 and 100 μM HCQ. Insets summarize the voltage clamping protocols. Right panels (ii): Mean concentration-response plots constructed for the maximum magnitude of each current ((n cells).

**Figure 3.**
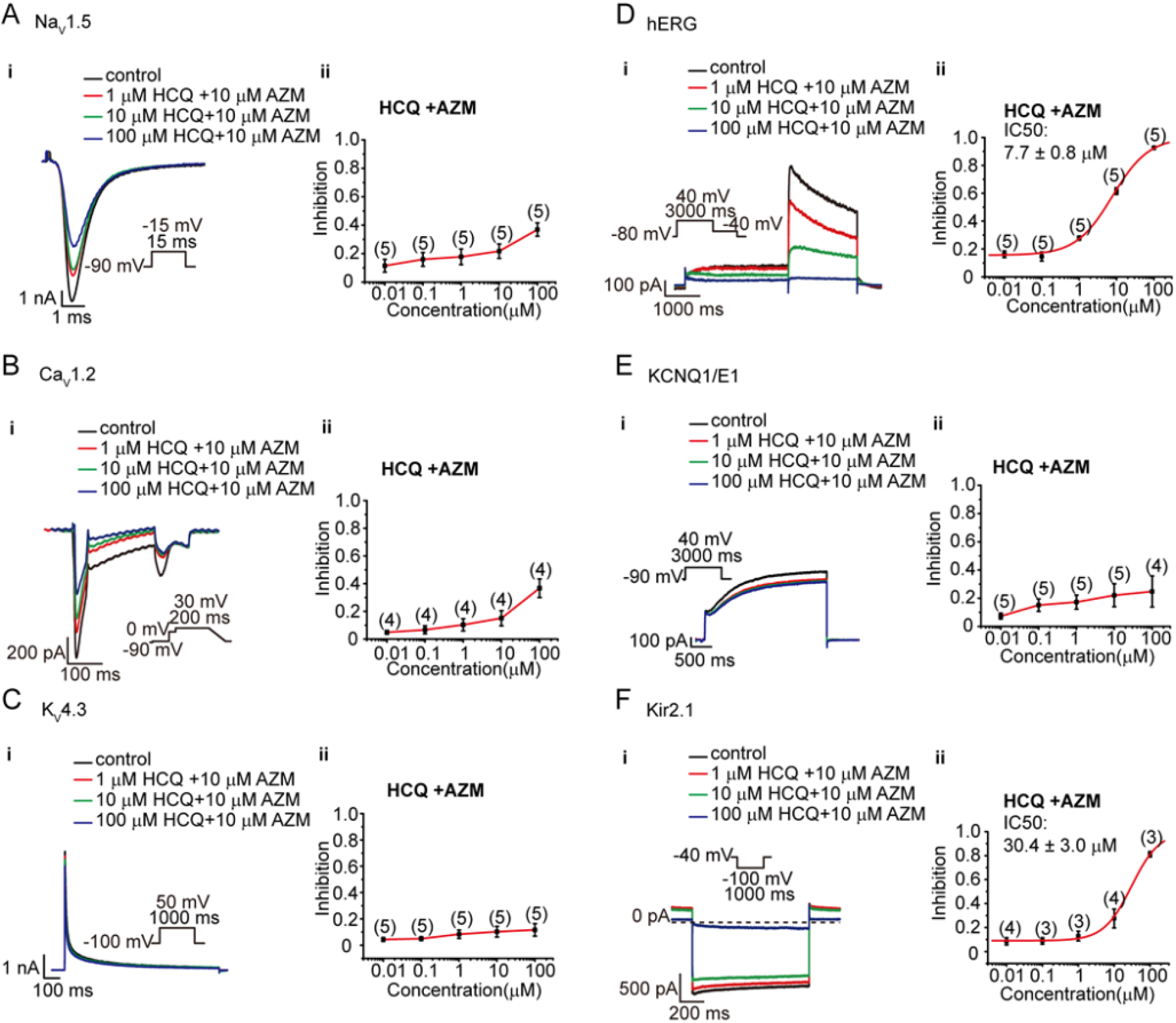
Effects of HCQ in combination with 10 μM AZM on ionic currents attributable to hNav1.5 (I_Na_, A), Cav1.2 (I_Ca,L_, B), Kv4.3 (I_to_, C), hERG (I_Kr_, D), KCNQ1/E1 (I_Ks_, E) and Kir2.1 (I_K1_, F). (A-F) Left panels (i): Typical currents in the presence of 0, 1, 10 and 100 μM HCQ with 10 μM AZM. Insets summarize the voltage clamping protocols. Right panels (ii): Mean concentration-response plots constructed for the maximum magnitude of each current (n cells).

Figure1 (left panels, i) exemplifies observed ionic currents in the presence of 0, 1, 10 and 100 μM HCQ. The insets summarize the related voltage clamping protocols, fully described in extended Methods in Supplementary data. Five concentrations ranging from 10^-8^ to 10^-4^ M were tested for each current examined in 3 to 6 cells at each concentration. Mean concentration-response plots (right panels, ii) used the maximum magnitudes of each current. Where applicable, the consequent alterations in such current magnitudes following pharmacological challenge were used to derive IC_50_ values.

Figure1 (right panels, ii) plots mean (±SEM) fractional reductions in current magnitude against HCQ concentration for I_Na_ (A), I_CaL_ (B), I_to_ (C), I_Ks_ (D), I_Kr_ (E) and I_K1_ (F). μM-HCQ inhibited both I_Kr_ tail and I_K1_ pulse currents with IC_50_s of 10±0.6 μM (D) and 34±5 μM (F), respectively, with more marked effects on I_Kr_, where the IC_50_ fell close to expected therapeutic levels in current Covid-19 treatment regimes (6), than I_K1_. In contrast, μM-HCQ has less effect on I_Na_ and I_CaL_ with IC_50_s of 113 ± 78 μM and 209 ± 94 μM respectively, Nevertheless, HCQ caused graded I_Na_ reductions through the entire HCQ concentration range typified in a 22±1.8% reduction at 1 μM. No significant inhibition of I_to_ and I_Ks_ by μM-HCQ was observed.

Figure 2 presents corresponding results obtained with AZM applied by itself similarly encompassing clinical therapeutic levels. A previously reported dose regime administering 500, 250 and 250 mg/day on each of three successive days gave a day 2 serum concentration of 0.22 μM(15)). However, the entire, 0-100 μM explored AZM concentration range yielded only minimal inhibitory effects. Even 10 μM and 100 μM AZM reduced I_Na_ (A), I_CaL_, (B) I_Ks_ (E) and I_Kr_ (D) by <17% and <35% respectively and produced negligible effects on I_K1_ (F) and no effect on I_to_ (C).

Finally, Figure 3 summarizes results of applying the same range of five HCQ concentrations displayed in Fig. 1 now combined with 10 μM AZM. Adding AZM did not significantly alter the effects of HCQ on I_Na_ (A), I_CaL_ (B), I_to_ (C), and I_Ks_ (E), but reduced the IC_50_ for I_Kr_ (IC_50_ = 7.7 ± 0.8 μM) and I_K1_ (30.4 ± 3.0 μM).

### Computational molecular docking determining the interaction between HCQ, AZM and hERG

The above findings particularly concerning pharmacological effects on hERG prompted computational molecular docking explorations for possible interactions between HCQ and AZM and a recent hERG homotetramer structure (PDB ID: 5va1; 1) (Fig.4). We used scoring docking visual inspection irrespective of the docking protocol. Figure 4 illustrates the HCQ-hERG interaction, derived using three separate rigid-receptor docking methods. HCQ interacts predominantly with Y652 from one or more subunits. Interactions with other residues including F656 but also S624, A653, and S660 occurred less frequently. Variability in observed docking positions could be attributed to the highly symmetric structure of the hERG channel pore constituting the binding site enabling many equivalent positions. In contrast, no interactions were demonstrated between AZM and hERG (data not shown).

**Figure 4.**
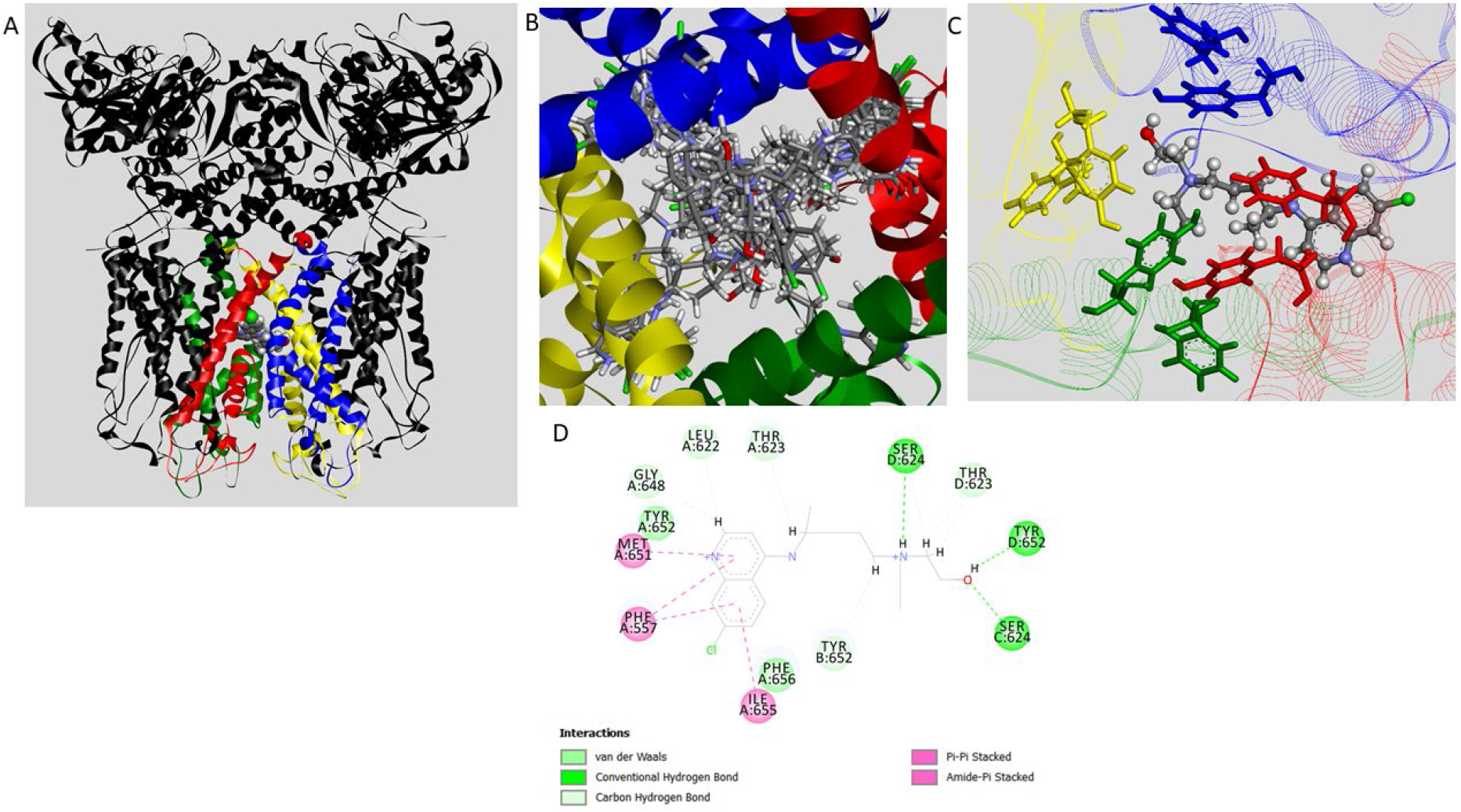
Computational docking studies. (A) HCQ (ball-and-stick model) docked in hERG structure. Y652 and F656 residues in stick representation colored accordingly to its parent subunit A, B, C or D (red, green, yellow, and blue, respectively). (B) 2-dimensional (2D) diagram depicting interaction types between presented HCQ pose and hERG. Diagram was prepared using Non-bond Interactions Monitor in Discovery Studio Visualizer (Dassault Systèmes, Vélizy-Villacoublay, France).

### Electrical mapping and electrocardiographic studies in *ex vivo* heart preparations

We then explored the effects of HCQ alone and combined with AZM in the whole heart on electrophysiological activation and recovery, their electrocardiographic correlates and the consequent Ca^2+^ signalling processes reflecting excitation-contraction coupling.

Figure 5A summarises sites of apical stimulation, and multi-electrode array (MEA) and ECG recording in the electrical mapping experiments. Representative left atrial (LA) and left ventricular (LV) isochronal conduction maps (B) obtained in the presence of 0 - 10 μM HCQ revealed a clear reduction of conduction velocity and increased conduction heterogeneities at 10 μM HCQ. Synergistic inhibitory effects on conduction velocity were observed when hearts were treated with the 10 μM HCQ in combination with 10 μM AZM. All these effects were reversed on drug washout (C(i)). 10 μM HCQ significantly decreased conduction velocities of excitation with further decreases in velocity with the 10 μM HCQ and 10 μM AZM combination (C(ii)).

**Figure 5.**
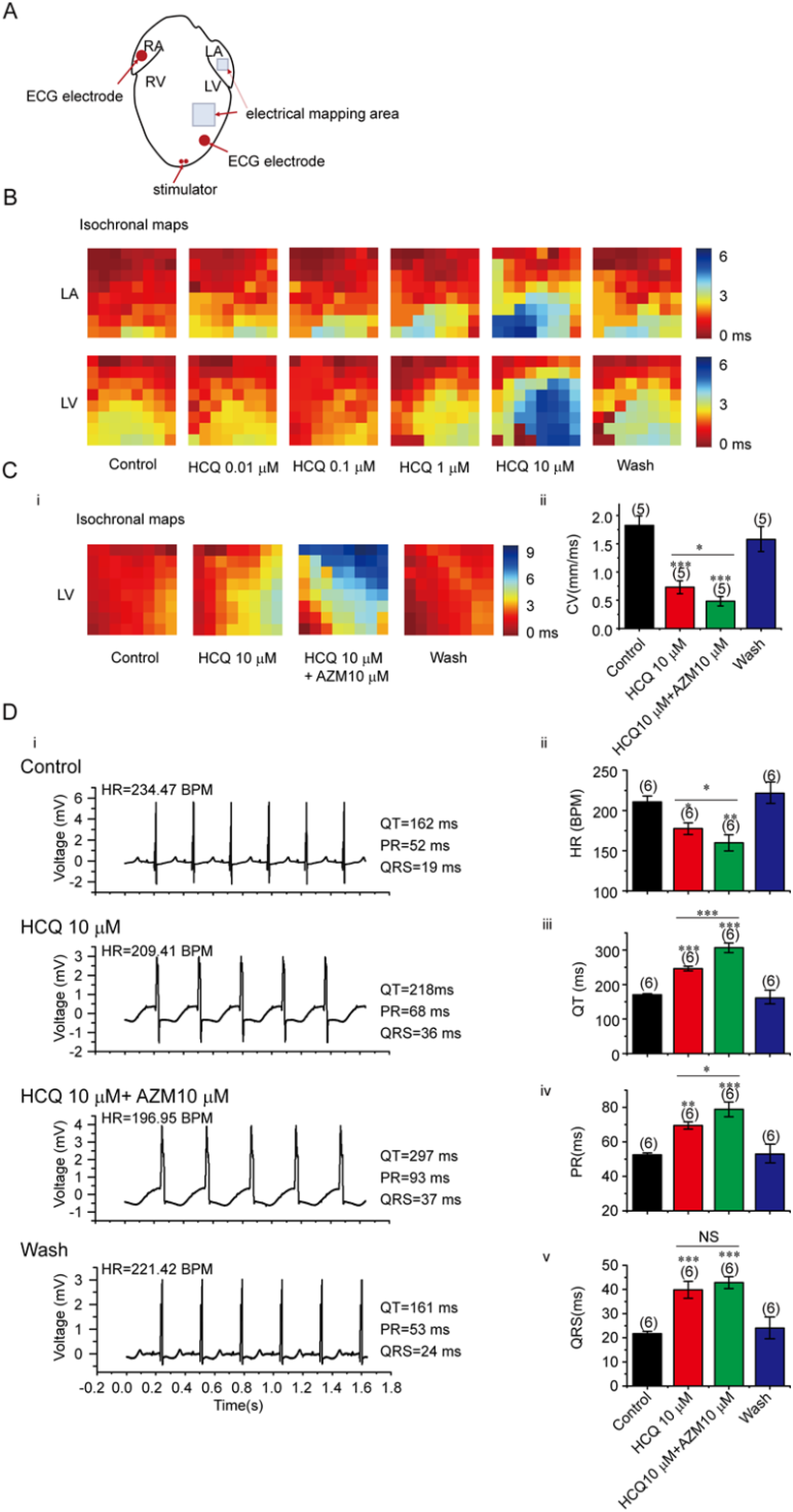
Multi-electrode array (MEA) mapping and electrocardiographic (ECG) studies in isolated perfused guinea-pig hearts following HCQ and AZM challenge. (A) Schematic summary of stimulation and MEA and ECG recording configuration. (B) Typical left atrial and left ventricular isochronal conduction maps obtained in the presence of 0, 0.01, 0.1, 1.0 and 10 μM HCQ. (C) (i) Successive LV isochronal maps before and following addition of 10 μM HCQ and 10 μM HCQ and 10 μM AZM in combination followed by drug washout. (ii) Quantified conduction velocities through these conditions. (D) ECG studies: (i) ECG records obtained before and following application of 10 μM HCQ, and 10 μM HCQ and 10 μM AZM combined, followed by drug washout. (ii)-(v): corresponding measured (ii) heart rates (HR), (iii) QT and (iv) PR intervals and (v) QRS durations (n hearts) * p<0.05, ** p<0.01, *** p<0.001

Quantification of the corresponding electrocardiogram (ECG) traces (D(i)) demonstrated that both 10 μM HCQ, and 10 μM HCQ in combination with 10 μM AZM, decreased heart rate (HR)(D(i)), and increased QT interval (D(iii)), PR interval (D(iv)) and QRS duration (D(v)). In particular, AZM when combined with HCQ augmented its effects on HR, and QT and PR interval. Together these results suggested that 10 μM HCQ, and 10 μM HCQ and 10 μM AZM combined, exerted synergistic inhibitory effects on heart rate and conduction, and prolonged cardiac repolarisation.

### Optical mapping studies in *ex vivo* heart preparations

We next investigated the effects of HCQ alone and in combination with AZM using optical fluorescence mapping in sinus-paced Langendorff-perfused hearts using the recording and ECG monitoring configurations illustrated in Figure 6A(i). Voltage mapping measurements of action potential initiation and conduction (Fig. 6A(ii)) demonstrated bradycardic effects (A(iii)), visible conduction heterogeneities and slowed conduction (A(ii)) at 10 μM HCQ, effects were augmented by combination with 10 μM AZM (Fig. 6A (iv)). Similarly, optical potentiometric mapping revealed that 10 μM HCQ prolonged action potential duration at 90% repolarisation (APD_90_) (B(i and ii)) with 10 μM HCQ, effects further prolonged by its combination with AZM (Fig.5B(iii)).

**Figure 6.**
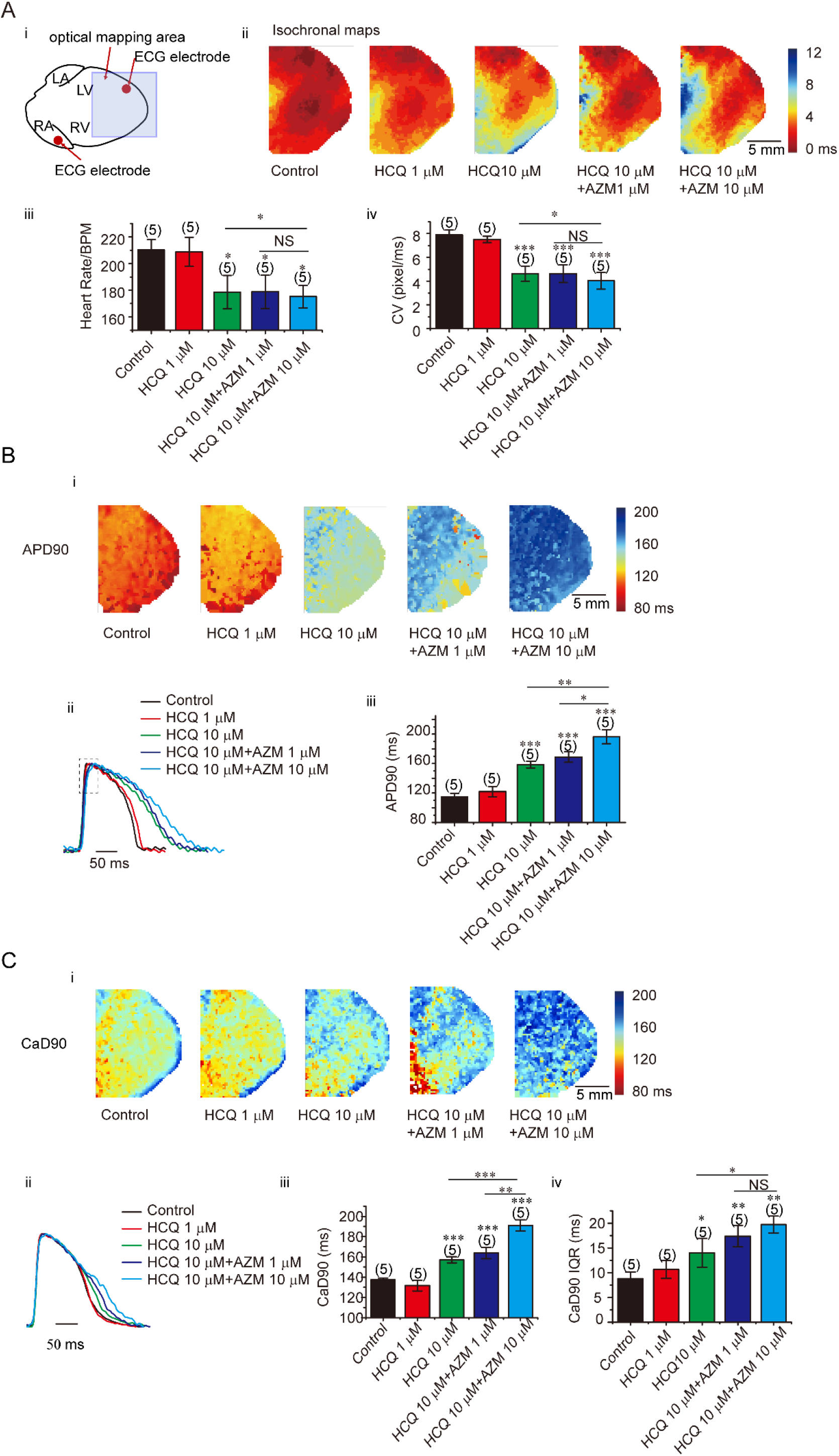
Voltage RH237 and Rhod-2 optical mapping in isolated perfused hearts. Comparisons of results before and following challenge by 1 μM and 10 μM HCQ, and 10 μM HCQ combined with either 1 or 10 μM AZM. (A, B) RH237 mapping: (A)(i) Optical mapping and ECG monitoring configuration (ii) Maps of action potential initiation and conduction; (iii) heart rates and (iv) conduction velocities. (B) Action potential time-courses and recoveries: (i) Maps of AP durations at 90% repolarization (APD_90_). (ii) Representative APs recorded from defined regions of interest. (iii) APD_90_s averaged over the field of view. (C) Rhod-2 AM (CaD) mapping of ventricular Ca^2+^ transients (CaT): (i) maps of CaD durations at 90% recovery (CaD_90_). (ii) Comparison of typical CaT transients averaged over defined regions of interest; (iii) CaD_90_ magnitudes; (iv) CaD_90_ dispersions (n hearts). NS not significant, * p<0.05, ** p<0.01, *** p<0.001

**Figure 7.**
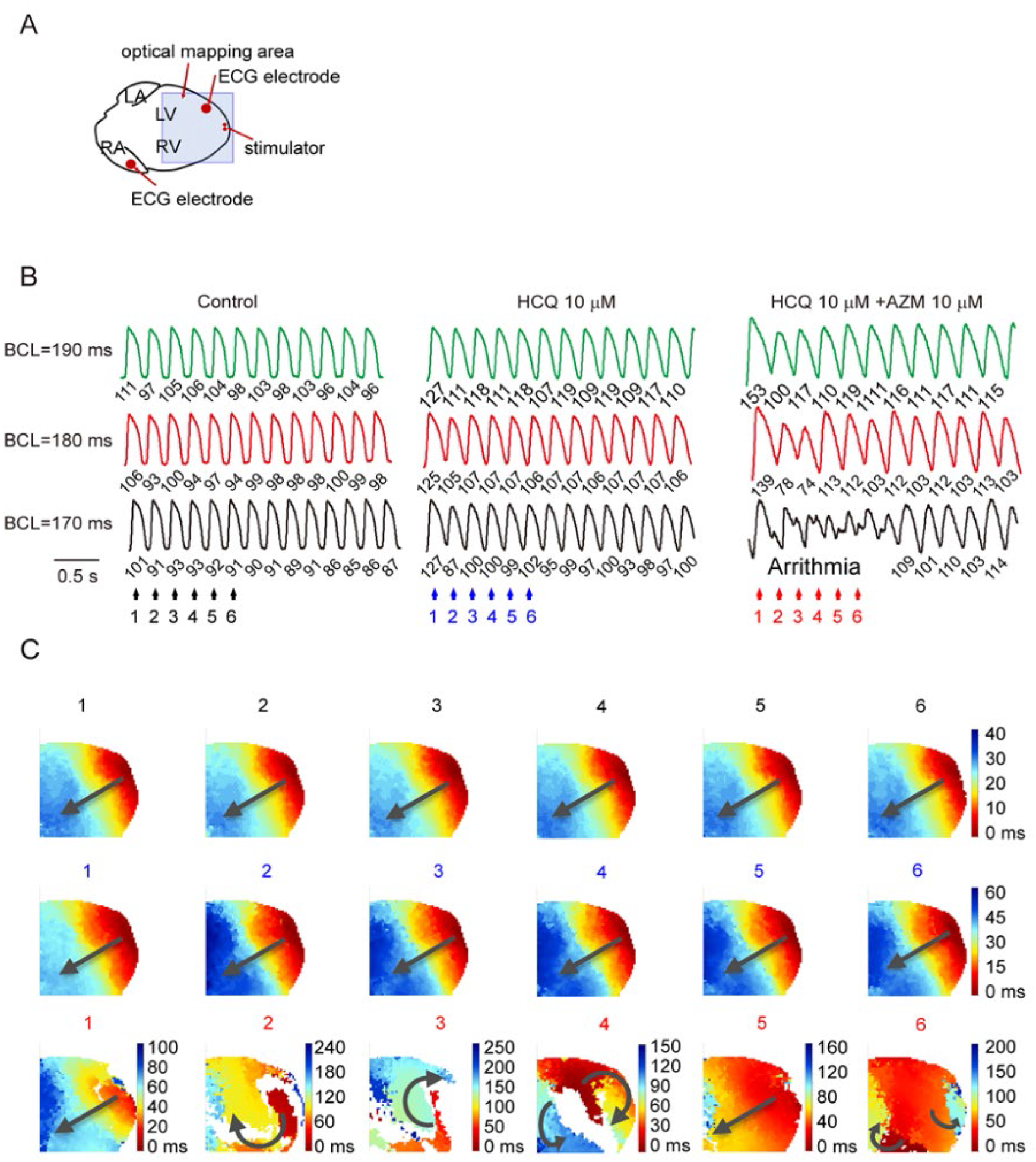
Voltage RH237 optical mapping of re-entry and ventricular arrhythmic tendency in isolated perfused hearts. (A) Recording and programmed stimulation configuration. (B) Optical action potential traces obtained under programmed pacing at 190 ms, 180 ms and 170 ms cycle lengths (CLs), before (control) and following addition of 10 μM HCM, and 10 μM HCM with added 10 μM AZM. (C) Corresponding voltage maps obtained at the numbered colour-coded timepoints (n hearts).

Ca^2+^ dye, Rhod-2 AM (CaD), mapping of ventricular calcium transient (CaT) characteristics demonstrated increased magnitudes of and dispersions in CaD durations at 90% recovery (CaD_90_) (Figure 6C(i)). The latter were attributed to changes late in the CaT records (C(ii)). The quantified CaD_90_ magnitudes (B(iii)) and dispersions (B(iv)) both increased with 10 but not 1.0 μM HCQ. The combination of 10 μM HCQ with either 1 or 10 μM AZM exerted significantly greater effects than 10 μM HCQ alone.

### Optical mapping assessments of arrhythmic tendency in *ex vivo* heart preparations

Finally, voltage dye optical mapping studies during ventricular pacing were used to assess susceptibility to arrhythmia induction before and after drug challenge in isolated perfused hearts (Fig.7A.B). Whilst 10 μM HCQ did not exert detectable arrhythmic effects at the cycle lengths (CL) used, a combination of HCQ and AZM produced AP alternans at 180 ms CL and episodes of non-sustained torsadogenic-like VT at 170 ms CL in 5 out of 5 hearts. The corresponding potentiometric maps obtained during the colour coded timepoints 1-6 are represented in C. At a CL of 170 ms, the conduction maps with HCQ+AZM demonstrated significant disruptions of the normally coherent propagation of excitation waves with evidence of excitation re-entry and wave break. Thus, the electrophysiological changes outlined above associated particularly with HCQ and AZM in combination were associated with a ventricular arrhythmic tendency.

### Computational simulations of HCQ and AZM on human ventricular action potentials

To translate the experimental data describing the multi-ion channel blocking effects of HCQ and/or AZM to human ventricular excitation, the ORd model was implemented to reconstruct the impacts of HCQ alone and combined with AZM, on human ventricular action potentials including APD_90_ and maximal upstroke velocities.

Figure 8A displays predicted action potentials under control conditions and in the presence of HCQ, and HCQ and AZM combined. It was shown that with the 10 μM HCQ alone, the APD was prolonged by 100 ms as compared to the control condition. With the combined action of HCQ and AZM, a synergistic action of the two drugs on APD prolongation was seen, which was manifested in a greater increase of APD90 of ~300 ms (Figure 8B). Such an APD prolongation can be attributable to the integrated action of reduced repolarisation potassium channel currents of I_Kr_ and I_K1_, and reduced depolarising currents of I_Na_, I_CaL_ and I_NaCa_. A small reduction in action potential maximum upstroke velocities was also observed by both HCQ and HCQ+AZM due to the effects on I_Na_.

**Figure 8.**
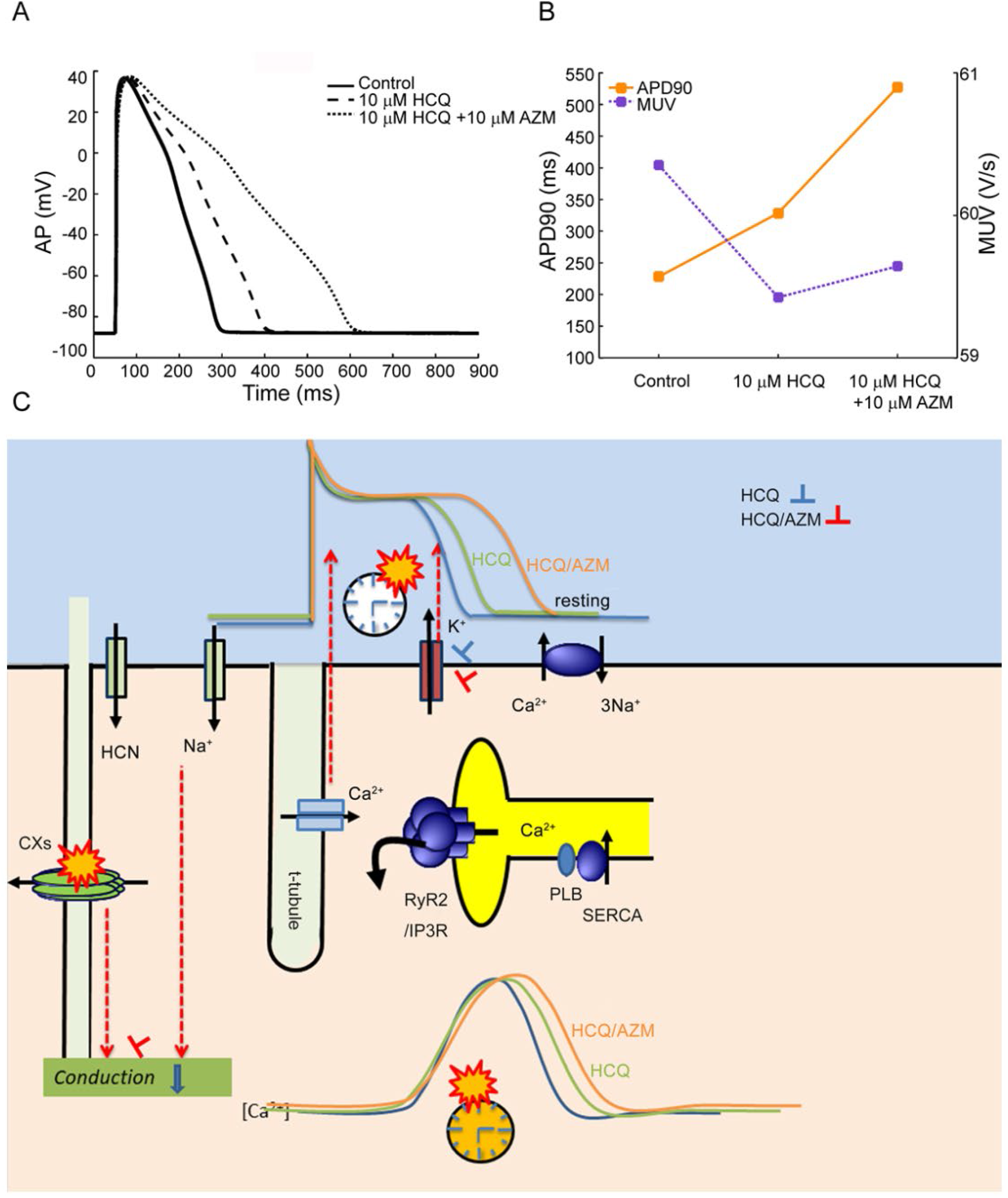
In silico modelling of action potentials before and following HCQ and combined HCQ and AZM challenge. (A) Simulated action potentials of human ventricular cell under control conditions and in the presence of HCQ (10 μM) and HCQ (10 μM) + AZM (1 μM) combined. HCQ and AZM when combined produced a synergistic action in prolonging action potential duration (APD_90_), as well as reducing the repolarization rate following the end of phase 0 depolarisation. (B) Effects of HCQ and HCQ + AZM combined on APD_90_ and the maximal upstroke velocity (MUV) of the AP. (C) Illustration of electrophysiological effects and possible mechanisms underlying the pro-arrhythmic effects of applications of HCQ and HCQ and AZM combined at the ion channel, cellular and tissue levels.

## DISCUSSION

The main findings of this study are: (1) Reported therapeutic levels of HCQ blocked multiple human cardiac ion channels, particular hERG (mediating I_Kr_) and Kir2.1 (I_K1_,) and, to a lesser degree, hNav1.5 (I_Na_). AZM potentiated the effects of HCQ on hERG and Kir2.1. (2) Computational molecular docking studies correspondingly predicted HCQ binding to hERG. Accordingly, (3) at the level of intact hearts, HCQ increased QT intervals as well as ventricular AP and Ca^2+^ transient durations and their heterogeneities. However, (4) HCQ further reduced heart rates, A-V and ventricular conduction as well as increasing conduction heterogeneities. All these effects were accentuated by combining HCQ challenge with AZM. Furthermore, (5) The HCQ+AZM combination, but not HCQ alone, then went on to increase susceptibilities to ventricular arrhythmic events consistent with significant clinical cardiac safety risks. (6) Human *in-silico* modelling of ion channel effects of HCQ and AZM recapitulated the altered APDs but not the reduced conduction. This mechanistic electrophysiological data complements ongoing multiple randomized trials evaluating clinical efficacy and cardiac safety profiles of these drugs as a treatment for Covid-19. The findings directly align with recent FDA guidelines concerning their use.

These successive lines of investigation were combined to reach an integrated view of HCQ and AZM action. This began with the approach recommended by the CiPA guidelines (12) in which we examined effects of HCQ and AZM on human cardiac ion channels responsible for the main inward (hNav1.5 (I_Na_) Cav1.2 (I_Ca,L_)) and outward (Kv4.3 (I_to_), hERG (I_Kr_), KCNQ1/E1 (I_Ks_) and Kir2.1 (I_K1_)) currents in the human ventricular myocardium. HCQ and AZM were studied both alone and in combination through a wide concentration range encompassing and extending beyond reported therapeutic levels. The latter typically fell around 1.15-4.5 μM(16) and 0.5-0.7 μM(17) for HCQ in SLE and rheumatoid arthritis patients on a daily 400 mg dose. In contrast, its dose used in Covid-19, 1.2 g/day, would give substantially higher therapeutic concentrations of ~10 μM(6). Blood AZM levels following successive daily 500 mg and two 250 mg doses were reported at 0.22 μM(15). HCQ both alone and combined with AZM synergistically inhibited I_Kr_ and I_K1_ with IC_50_s at the high end of reported therapeutic levels, with some minor inhibition of I_Na_ in this range.

Consistent with these experimental findings, computational molecular docking simulations utilizing a recent hERG homotetramer structure (PDB ID: 5va1; 1) demonstrate potential HCQ binding. This is predominantly with Y652 from one or more hERG subunits. Interactions with other residues including F656 but also S624, A653, and S660 occurred less frequently. There was no predicted AZM-hERG binding in the model although this does not discount it occurring *in-vivo*.

Others have also explored the electrophysiological effects of HCQ or AZM on some cardiac ion channels but did so at three times the clinically relevant concentration and above (15, 18). Furthermore, the present study systematically evaluates HCQ and AZM in combination for the first time. Furthermore, the only report on electrophysiological effects of HCQ specifically studied sinoatrial cells, demonstrating significant reductions in hyperpolarization-activated currents (If), with reduced I_CaL_ and I_Kr_ observed at 3 μM (18). Nevertheless, this HCQ action on If explains its bradycardic effects observed here, further illustrating its broad cardiac electrophysiological actions. Such a bradycardic medication in the presence of a hypotension accompanying sepsis or SARS could potentially exacerbate a septic shock.

At the whole heart level, our optical mapping studies demonstrate that HCQ correspondingly prolongs ventricular APD as well as CaD, to extents further exacerbated by combination with AZM. This fulfils expectations from its inhibition of I_Kr_ and I_K1_ and translates to the observed electrocardiographic QT prolongations. It is interesting to note that even prior to the Covid-19 pandemic, series of case reports have led to both HCQ and AZM being listed as definite causes of prolonged QT interval and risk of torsade de pointes for patients with long QT syndrome on *crediblemeds.org* (19).

Prolonged APD predisposes to early afterdepolarisations (EADs) during the plateau phase of the ventricular action potential, classically attributed to I_CaL_ reactivation (20). EADs are also more common under conditions of intracellular Ca^2+^ overload, including increased sympathetic drive: I_CaL_ can be enhanced by high intracellular Ca^2+^ levels and by L-type Ca^2+^ channel phosphorylation by CaMKII(21). EADs may potentially trigger ectopic beats and initiate arrhythmic events given suitable substrate.

In addition to these recovery phenomena, we also demonstrate for the first time that HCQ reduces ventricular conduction on multi-electrode arrays and optical mapping recording, and compromise atrioventricular and intraventricular conduction, reflected in prolonged PR intervals and QRS durations, to extents accentuated by further addition of AZM. Such findings are consistent with Na channel block (22) of the kind exemplified clinically by the CAST trials (23) and/or compromised gap junction function(24). Together, the combination demonstrated here of prolonged and increased dispersions of repolarisation and calcium handling combined with slowed conduction would offer a suitable substrate for sustained re-entrant arrhythmia. Thus, the combined exposure to HCQ and AZM, but not HCQ alone provoked alternans, ventricular tachycardic episodes, and re-entrant excitation following progressively accelerated cardiac pacing.

Finally, a validated human ventricular myocyte mathematical model integrated the observed ion channel properties with the effects on action potential and intracellular calcium handling characteristics in intact hearts. It separated whole heart electrophysiological features that were explicable in terms of the single channel properties and those attributable to altered calcium homeostasis. The modelling reproduced effects of surface ion channel function and their interactions forming a ‘membrane clock’ underlying the measurable electrical activity (Fig. 8C)(14). It reproduced the action potential prolongation and consequent prolonged QT intervals with HCQ and the synergistic effects of combining HCQ with AZM, attributing these to their combined effects on I_Kr_ and I_K1_. These latter events have been described as feeding forward into a ‘Ca^2+^ clock’ driving excitation-contraction coupling but whose events exert feedback effects modifying action potential conduction, recovery, and post-recovery stability, exemplified by Ca^2+^ feedback effects on Na^+^ channel and gap junction function (Fig. 8C), as well as longer-term effects on channel expression(13, 25). The modelling predicted the increased calcium transient magnitudes and durations particularly with HCQ+AZM (Fig 8C), but only small changes in maximum upstroke velocity attributable to Na channel blockade. The observed conduction slowing nevertheless could arise from a Ca^2+^-dependent gap junction uncoupling.

These results together provide an integrated basis for the electrophysiological and pharmacological effects of HCQ, AZM and their combination. They thereby underpin recent FDA guidelines cautioning against combined HCQ/AZM administration for treatment of Covid-19 on grounds of cardiac safety. Admittedly, physiological though not pro-arrhythmic effects occurred with HCQ administered alone at the ~10 μM concentrations associated with Covid-19 therapy and not at the lower ~1.0 μM HCQ levels associated with therapy for rheumatoid arthritis and SLE. HCQ alone at therapeutic doses thus did not increase incidences of experimental ventricular arrhythmias in healthy hearts. Nevertheless, safety threshold doses could be reduced by accompanying cardiac injury arising from viral toxicity, sepsis, hypoxia-related myocyte injury, and immune-mediated cytokine storm in Covid-19. Furthermore, AZM potentiated the HCQ toxicity, now producing arrhythmic outcomes. The present findings thus strongly indicate a desirability for monitoring electrocardiographic, including QT, changes and cardiac injury during 4-aminoquinoline-macrolide therapy for Covid-19 patients.

## METHODS

### Study Approval

All procedures were approved by The Institutional Animals Ethics Committees at Henan University, Kaifeng, China under and the national guidelines under which the institution operates, and also conform with the NIH Guide for the Care and Use of Laboratory Animals. Guinea pigs (male; 300-350 g) were provided by The Experimental Animal Center of the Medical College of Xi'an Jiaotong University and maintained in a pathogen−free facility at Henan University with *ad libitum* access to food and water. Animals were humanely sacrificed by intraperitoneal injection with 3% pentobarbital sodium (50 mg/kg).

### Single-cell electrophysiological recordings

Core human cardiac channels were heterologously expressed in recombinant HEK293 or CHO cell lines and studied using conventional patch clamp methods. Stable cell lines expressing Nav1.5 and Cav1.2/β2/α2/δ1, hERG, Kir2.1, Kv4.3, KCNQ1/E1 were used. All currents were recorded using a MultiClamp 700A amplifier, 1550A digitizer and pCLAMP 10.6 software (Molecular devices, USA). Patch electrodes with resistances of 2-5 MΩ were pulled with a P-97 micropipette puller (Sutter Instruments, USA). The conventional whole-cell recordings were obtained by manual suction for hERG, Kir2.1, Kv4.3 and Nav1.5 studies. Perforated whole-cell recordings of KCNQ1/E1 and Cav1.2 were obtained following perforation with 200 μg/ml Amphotericin included in the pipette solution. After whole-cell mode were obtained, cells were used in voltage clamp mode for the current recordings. The external solution contained (mM): NaCl 137; MgCl_2_ 1; KCl 4; glucose 10; HEPES 10; CaCl_2_ 1.8, pH adjusted to 7.4 with NaOH. The K^+^-containing pipette solution contained (mM): KCl 40; KF 100; MgCl_2_ 2; EGTA 5; HEPES 10, pH adjusted to 7.2 with KOH. The sodium and calcium current pipette solution contained (mM): CsCl 65; CsF 75; MgCl_2_ 2.5; EGTA 5; HEPES 10, pH adjusted to 7.3 with CsOH.

### Molecular modelling and docking studies

The experimental structure of the open hERG channel (PDB ID: 5va1; 1) was selected as a receptor for molecular docking studies. The structure was prepared for docking using Protein Preparation Wizard using default settings including modelling of missing loops using Prime. The loops spanning residues 433-448, 578-582, and 588-602 in each subunit were successfully rebuilt. The centroid of residues Y652 and F656 from every subunit was defined as the centre of the docking binding site. The HCQ structure was obtained from PubChem and prepared for docking using LigPrep. Three separate rigid-receptor docking methods were used including: (i) Glide SP docking, (ii) FlexX docking followed by HYDE re-scoring, and (iii) AutoDock Vina docking with each set to output 20 poses per LigPrep HCQ state. Poses from Glide, FlexX, and AutoDock Vina were also re-scored using RF-Score-VS.

### Langendorff-perfused isolated hearts

Guinea pigs (male; 300-350 g) were humanely killed by intraperitoneal injection with pentobarbital sodium (50 mg/kg). Hearts rapidly excised after thoracotomy were mounted onto a Langendorff perfusion system and perfused with a modified Tyrode’s solution (119 NaCl, 25 NaHCO_3_, 4 KCl, 1.2 KH_2_PO_4_, 1 MgCl_2_, 1.8 CaCl_2_.2H_2_O, and 10 D-glucose (mM) equilibrated with 5% CO_2_ and 95% O_2_) with the flow rate of 8 ml/min at 37°C. Hearts were perfused and monitored for stability for 20 mins before experimental procedures commenced.

### Electrical mapping of *ex vivo* heart preparations

A multi-electrical array (MEA) mapping system (EMS64-USB-1003, MappingLab Ltd., UK) was employed to record extracellular potentials (ECP) from the epicardium. Two 64 channel MEAs (MappingLab Ltd., UK) were employed for left atrial and left ventricular simultaneous recordings. ECP signals were recorded at a 10 kHz sampling using synchronized 64 channel EMS64-USB-1003 recording systems and recorded using EMapRecord 5.0 software (MapingLab Ltd., UK) and stored in a computer for subsequent off-line analysis. Electrocardiograms (ECG) were continuously recorded by two ECG electrodes (MappingLab Ltd., UK) positioned on the right atrium and left ventricle respectively.

The epicardial isochronal activation map during sinus rhythm or imposed pacing was plotted as time differences between the earliest activation site and the individual activation site on each channel. Activation times were calculated as the point of maximal negative slope of activation waveforms. The conduction velocity (CV) was calculated from the difference in timing and the known distance between the recording points using EMapScope 5.0 software (MappingLab Ltd.).

### Optical mapping of *ex vivo* heart preparations

After the Langendorff-perfused hearts reached steady state, contraction artefacts were mimimised using blebbistatin (10 μM). Dye loading was aided by pre-perfusion with pluronic F127 (20 % w/v in DMSO). Rh237 (1μg/ml) and Rhod2-AM (1μg/ml) were perfused to enable simultaneous membrane potential and Ca^2+^ measurements at 37 °C.

Two 530 nm LEDs (LEDC-2001, MappingLab Ltd) were used to illuminate the heart after their emissions were bandpass filtered (wavelengths 530±20 nm) to minimize stray excitation light reaching the dyes. The fluorescence light was passed through a 550 nm long-pass filter and then a dichroic mirror with a cut off of 638 nm. Fluorescence light with wavelengths above 638 nm was passed through a 700 nm long-pass filter and then imaged by the camera for recording voltage signals. Fluorescence light below 638 nm was passed through a bandpass filter (585±40 nm) then imaged by the camera for recording calcium signals (OMS-PCIE-2002, MappingLab Ltd). The raw spatial resolution was 128-by-128 pixels, the total mapping area was 16×16 mm and the temporal resolution was 900 frames/second.

The cameras of the optical mapping system, LED lights and stimulator and ECG recording were simultaneously driven by an 8 channel TTL analog-digital converter and OMapRecord 4.0 software (MappingLab Ltd.). For the analysis of optical mapping signals and generation of isochronal maps, data were semi-automatically processed using OMapScope 5.0 software (MappingLab Ltd). In this experimental configuration, stable recordings for voltage (*V*_m_) and intracellular Ca^2+^ (CaT) signals could be obtained for >2 hours in preliminary experiments assessing system stability.

### Arrhythmia induction protocol

Stimuli were generated by an isolated constant voltage/current stimulator (VCS3001, MappingLab Ltd., UK). They were delivered with a platinum electrode onto the epicardial apex at an amplitude 1.5 × the diastolic voltage threshold and a 2 ms pulse width. The optical mapping was performed in conjunction with an “alternans mapping” protocol. The heart was consecutively paced at basic cycle lengths (BCL) from 210 with 10 ms decrement until 170 ms. The heart was paced 50 times with 2 ms duration pulses and the optical mapping recording was performed at the last 2 second in each episode. This protocol was performed both before and after drugs were delivered.

### In silico model reconstruction

The O'Hara-Rudy (ORd) human ventricular model(26) was modified to incorporate the actions on multi-channel block of HCQ, including the dose-dependent block of I_Na_,, I_CaL_,, I_to_, I_Kr_, I_Ks_ I_K1_ and the Na^+^-Ca^2+^ exchange current, I_NCX_. This model was selected as it was developed from and validated against experimental data from human ventricular cells and found suitable for studying actions of drugs. The simulations evoked action potentials by a supra-threshold stimulus in control runs and following application of HCQ, and combined application of HCQ and AZM. Action potential time courses characterized by AP durations at 90% repolarisation (APD_90_) and maximal upstroke velocity (MUP) and the underlying ion channel currents targeted by the drugs were recorded for analysis.

### Statistical analysis

All patch clamp recorded data were analyzed using Clampfit 10.6 (Molecular Devices, USA), OriginPro 8.0 (Origin Lab, USA) and Adobe Illustrator 10 (Adobe, USA). The concentration-response curve was fitted by the logistic equation *y*= *A*_2_+(*A*_1_−*A*_2_)/[1+(*x*/*x*_0_)^*p*^], where *x* is the drug concentration, and *p* is the Hill coefficient. All data are expressed as the means ± SEM. One-way ANOVA followed by multiple-comparison test was used to evaluate multiple test treatments. A value of p<0.05 was considered to be statistically significant. IC_50_ denotes the concentration determined for half-maximal inhibitory effects. In the figures, the designations for p values are: *p<0.05, **P<0.01 and ***P<0.001 respectively.

## Author contributions

GH, CL, AT, DP, CLHH, HZ and ML contributed to the study design

GW, XT, GH, YN, LW, XJ, GH contributed to the electrophysiological data acquisition.

GW, YD, GH, YX, LW, DL and NH contributed to the data analysis and visualization.

HF, KZ, HZ contributed to the action potential computer simulation.

PM and ML contributed to the molecular docking computer simulation.

LW, DP, ML and GH contributed to the funding and resource acquisition.

ML and GH contributed to the supervision.

All authors contributed to the writing.

